# Analysis of human BBS protein homologues in insects support alternative non-ciliary functions

**DOI:** 10.1101/2023.07.28.550953

**Authors:** Alexander Ewerling, Isa Graebling, Anna Wierczeiko, Elisa Kotzurek, Susanne Foitzik, Susanne Gerber, Thomas J. Colgan, Helen May-Simera

## Abstract

Cilia and flagella were one of the characteristic traits of the last eukaryotic common ancestor and as such, are highly conserved among eukaryotes. Their proteomic makeup is consequently remarkably similar throughout all eukaryotic lineages. Recently, one subgroup of ciliary transport proteins in mammalian cells, the Bardet-Biedl Syndrome (BBS) proteins, was shown to have the ability to traverse the nuclear envelope, and to engage in protein-protein-interactions that modulate gene expression, signalling cascades, and cell homeostasis. Insects have been critically understudied in cilia biology because of their highly specialised cilia being localised on only a small subset of cell types. In this study, we present evidence that the BBSome, a hetero-octameric ciliary transport complex of BBS proteins, is largely conserved in multiple insect lineages. Using the honeybee *Apis mellifera* as a study system to explore BBS-associated gene expression, our analyses suggest that not all BBSome-associated genes are expressed equally, indicating possible non-ciliary functions. We also demonstrate that the expression of individual BBS proteins varies significantly between the tissues of queens and males in *A. mellifera*, especially in neuronal tissue. This result raises the question of what role BBS proteins play in these tissues and whether they are involved in gene regulation in insects. The potential gene regulatory function of BBS proteins should be explored in other eukaryotes due to their high degree of conservation.

## INTRODUCTION

Cilia are tiny hair-like microtubule-based organelles extending from the surface of most eukaryotic cells. These ancient organelles have a conserved structure, function, and proteome across eukaryotes (1,2). They can be structurally divided into the basal body, which resides in the cytoplasm, the transition zone, the membrane-sheathed axoneme, and the ciliary tip (3) (Fig. 1A). While single-celled eukaryotes generally display motile cilia facilitating locomotion and sensory perception of the environment (2,4–6), immotile (or “primary”) cilia are mostly found as single copies per cell in metazoan cell types (7–9). Acting as a complex signalling centre, primary cilia are essential for several biological processes ranging from chemo- and mechanosensation to transduction of numerous signalling cascades. Defects in cilia can cause multiple multisystemic diseases, referred to as ciliopathies, which show a wide variety of partly overlapping phenotypes (10–12). One of these diseases, Bardet-Biedl Syndrome (BBS) (13,14), represents a genetically heterogeneous inherited disorder that is considered the archetypical ciliopathy since patients exhibit virtually all symptoms associated with ciliary dysfunction. These include retinal degeneration, kidney disease, obesity, and mental retardation. BBS is caused by pathogenic mutations in genes encoding BBS proteins, which are all involved in facilitating ciliary trafficking which underlies ciliary signalling (15–18) (Fig. 1A).

**Fig. 1:**
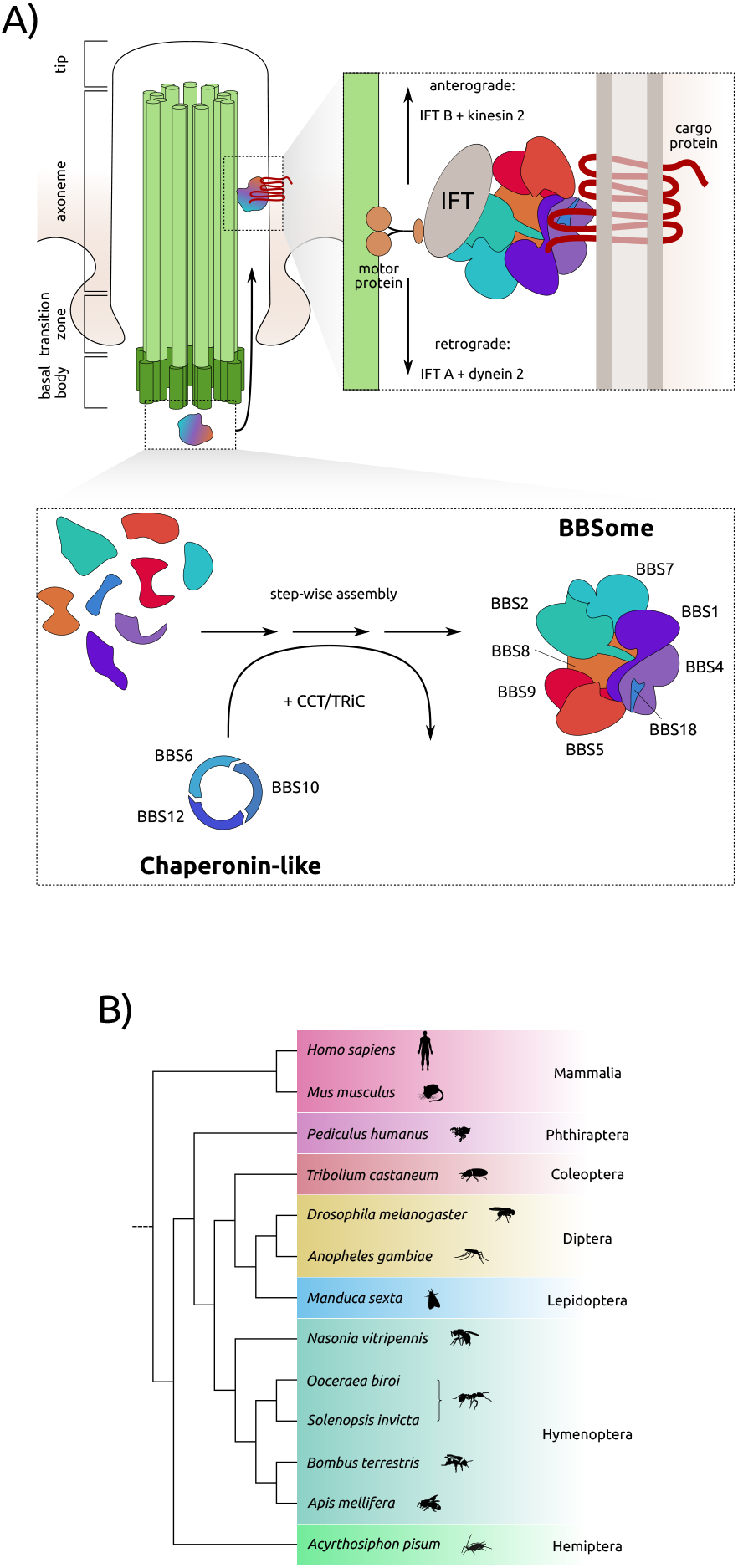
Ciliary functions are conserved in eukaryotes. A) BBS proteins can be found at the ciliary basal body where the BBSome is assembled with chaperonin-like BBS proteins in concert with CCT/TRiC proteins. Ciliary cargo proteins are then transported along the axoneme with the help of IFT particles and motor proteins (adapted from (18,37,58)). B) Although highly specialised, cilia can be found in insect neurons and sperm flagella. For orthologue search we assembled proteomes from several insect clades, and human and mouse as reference/outgroup.

Three main processes interact to make signalling pathways within cilia possible: trafficking of proteins to cilia from the cytoplasm, selective passage at the base of the cilium (the so-called ‘transition zone’), and specific intraflagellar transport (IFT) along the ciliary axoneme (15,19,20). The BBSome, a hetero-octameric complex consisting of the BBS proteins BBS1, BBS2, BBS4, BBS5, BBS7, BBS8, BBS9 and BBS18 (17,18,21), is considered an adaptor that couples ciliary cargo proteins to motor proteins and IFT complexes. In addition to the transport of vesicles to cilia, the BBSome is required to promote the retrieval and export of specific transmembrane proteins from the cilium (16,19,22–24). Therefore, the BBSome plays an important role as a key regulator of cilia composition. A second characteristic group of BBS proteins, so-called chaperonin-like BBS proteins, is comprised of BBS6, BBS10 and BBS12, which show structural homology to the chaperonin-containing t-complex protein 1 (CCT) family of group II chaperonins (25). It was previously demonstrated that these proteins form a hetero-oligomeric complex with other CCT proteins, and that this process plays a key role in the regulation of BBSome assembly (26,27). Cilia structure and function are highly dependent on specific proteins localising to the cilium and given that BBS proteins are required for this, the function of cilia directly depends on their functionality. In addition to the important role BBS and other ciliary proteins play in cilia, they have also been described to function in other cellular processes, such as cell cycle regulation (28) as well as regulation of gene expression by interacting with transcription factors and RNA polymerases (29) (for reviews, see (30–32)). Besides their ciliary functions, the biochemistry and modes of action of BBS proteins outside of cilia are largely unknown.

One approach to further elucidate molecular mechanisms of ciliary proteins in non-ciliary functions is from an evolutionary perspective. Cilia are present in most eukaryotes; however, they were lost independently on multiple occasions in distantly related taxa (33). Despite the absence of a classical cilia, genes coding for putative BBS proteins were shown to still be present in some of the genomes of these organisms. A prime example is the parasite *Toxoplasma gondii* where cilia assembly occurs independently of the BBSome. A single putative BBS5 gene homologue is still actively expressed in a non-flagellate state of the parasite’s life cycle (34), suggestive that the BBS-like protein does not perform cilia-associated functions and may have an alternative role. These non-ciliary functions may be crucial for an organism’s gene regulation as interactions with transcription factors and RNA polymerases have been identified (29). Consequently, these proteins might be essential beyond their ancestral ciliary functions. It is therefore of great interest to study the evolutionary patterns of putative BBS homologues and their expression in organisms that lack classical cilia. Indeed, insects possess those cilia in neurons and spermatozoa exclusively (35,36), and thus offer a unique opportunity to identify possible alternative functions of those genes through the determination of their expression outside of cells and tissues associated with the cilia.

As a preliminary step towards identifying possible alternative functions, we first determined whether there are homologues of human BBS proteins in representative members of diverse insect orders (Fig. 1B). More specifically, our study focussed on the BBS proteins that form part of the BBSome, as well as the genes encoding the chaperonin-like BBS proteins. The BBSome is ancestrally conserved in ciliated species and was predicted to be present in the last eukaryotic common ancestor (LECA) while the chaperonin-like BBS proteins are restricted to the unikont lineage (which includes animals and fungi) and the basal eukaryote *Malawimonas jakobiformis* (37), but are mostly found in metazoa. As mere presence or absence of protein homologues is a poor indicator of functionality, we also screened transcriptomic data of the well-studied honeybee *Apis mellifera* for expression of BBS proteins. Honeybees represent one of the most important pollinators, essential for the maintenance of agricultural yields and wildflower diversity (38). The genome was one of the earliest insect genomes to be sequenced (39) and assembled with the species having a wide range of transcriptomic and proteomic resources available (40,41). From an evolutionary perspective, honeybees are also part of the social Hymenoptera whereby caste differentiation has evolved within the female sex resulting in the expression of morphologically, behaviourally, and physiologically distinct phenotypes (42). Using publicly available datasets, we investigated sex- and tissue-specific expression differences to elucidate potential novel roles for BBS proteins in insects.

## RESULTS AND DISCUSSION

### Homology-based search reveals conservation of BBSome-associated proteins across insects

Our aim in this study was to investigate the occurrence and distribution of BBS proteins across highly divergent insect lineages. We started by studying the transcriptomes and genomes of different taxa and, in a second step, took a closer look at the proteomes of some insect species. To achieve this, we compiled a broad spectrum of predicted insect proteomes (n = 11 species across six orders; Fig. 1B). Divergence at the genome level is often so high that it is difficult to identify homologous genes with a high degree of certainty. However, since the protein sequence is much more conserved, protein-based analyses allow a closer look at the occurrence of homologous BBS proteins even in distant taxa.

In *Drosophila melanogaster*, cilia and flagella are only found on sperm cells and neurons (36). They still possess a functioning, relatively intact BBSome, albeit in a reduced form, with BBS2 and BBS7 missing from the otherwise complete complex found in other metazoa (37,43,44). We therefore investigated if a partial reduction (or even complete loss) is common across other insect taxa, or if the loss of single components identified in *D. melanogaster* is the onset of a taxonomically-restricted gradual loss of cilia. We screened the different insect species’ genome and transcriptome assemblies to compare conservation on nucleotide and peptide levels. Generally, over large evolutionary distances, gene-based orthology searches are only a weak indicator of presence or absence of a given gene, as codon bias might heavily skew the analysis, whilst proteins are highly conserved on an amino acid level, but not necessarily on nucleotide level. When BLAST-searching insect genomes with human nucleotide seeds we found a high rate of false-positive, and that only strict E-value cut-offs result in true hits (Fig. 2A). The case is similar when searching for transcribed BBS genes: almost none of the BBS genes appear to be transcribed in queried species, apart from BBS8. In addition to its role in ciliary trafficking, BBS8 has been shown to play a crucial role in centrosome stability and cell division (45–47). This suggests that: 1) the putative non-ciliary functions of BBS8 are independent of tissue, as inferred transcripts cannot be confidently assigned to a specific tissue of a given species; and 2) when searching for non-ciliary functions of BBS proteins, tissue-dependent expression plays a pivotal role in detectability. Considering the first point, it is interesting to find BBS4, BBS5, and BBS7 homologues also being expressed in some insect species. Previous studies showed a high probability for the proteins to be localised to the nucleus at some point, and also showed a nuclear localisation for the human orthologues (29,37). This merits further investigation into where these proteins might be expressed, especially in respect to tissue- and sex-dependent differences. As these gene products are most commonly found in the transcriptomic datasets across different species, the underlying basis for this conserved expression might be a conserved, non-ciliary function.

**Fig. 2:**
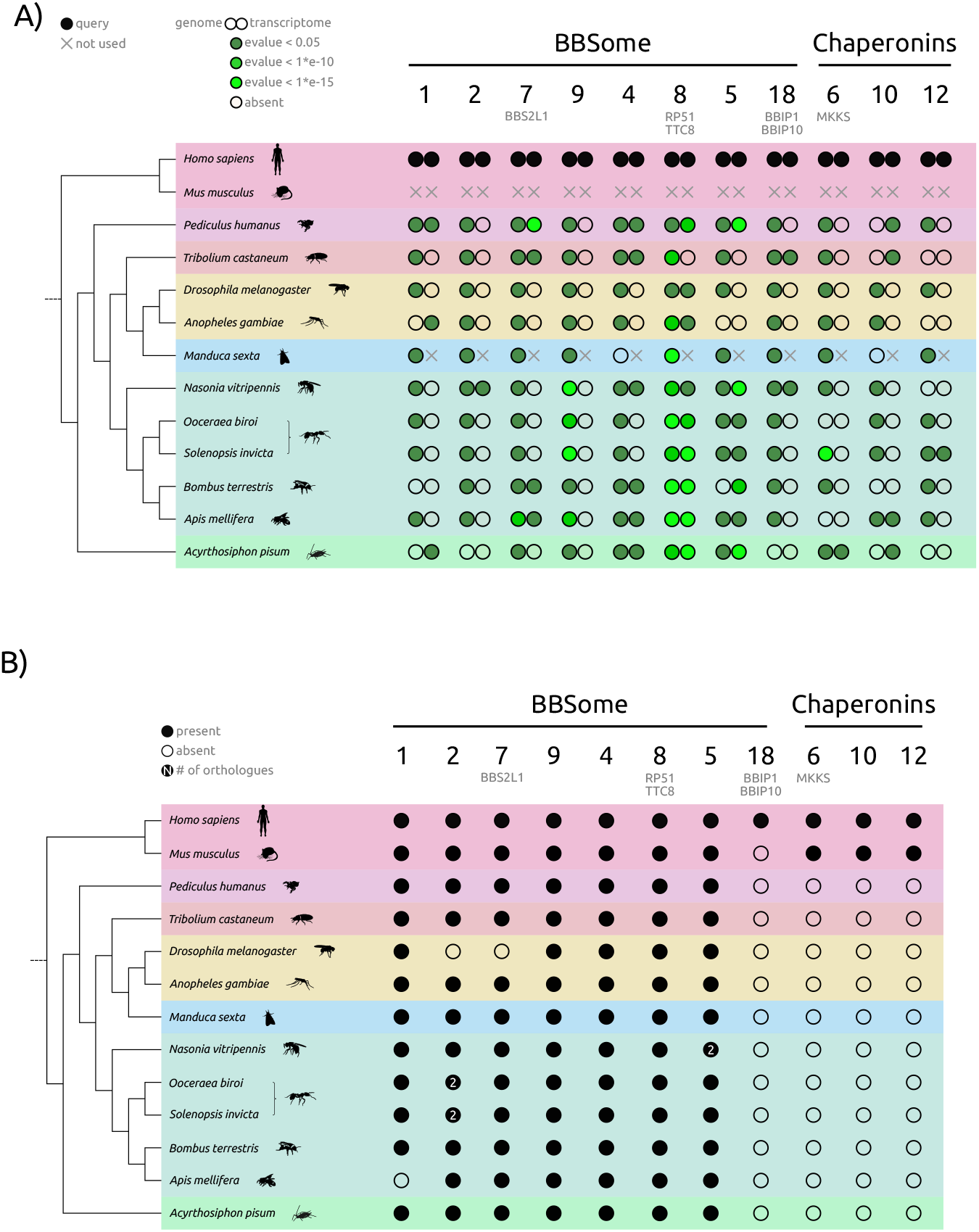
Evolutionary conservation of core BBSome genes across insects. A) Proteomic search has uncovered orthologues to most human BBSome proteins. BBS18 and chaperonin-like proteins BBS6, BBS10 and BBS12 are however restricted to humans in this analysis. B) While genome searches generally reflect more the proteomic searches, they are easily hitting false-positives (hits for almost all chaperonin-like BBS proteins).

To compare genome- and transcriptome-based searches with protein homologue searches, we compiled corresponding predicted proteomes from NCBI to conduct an all-versus-all BLAST-based homology search using OrthoFinder (48). We found that homologues of genes encoding BBS proteins that build the BBSome in humans (BBS1, BBS2, BBS7, BBS9, BBS4, BBS8, BBS5, BBS18) are present in almost all insect species, with the exception of BBS18 (Fig. 2B, Supp. Table S1). None of the queried species had homologues for chaperonin-like BBS proteins BBS6, BBS10 or BBS12. This lack of detection may be due to loss of genes in insects, gain of genes in mammals, or divergence of homologues beyond the point of detection. As cilia are built in highly specialised cell types in insects, there might not be a need for specialised CCT-derived BBS proteins in these cells to aid the folding of the BBSome. Overall, the high degree of conservation of the BBSome is reflected in most insect lineages, with *D. melanogaster* being a notable exception. BBS1-like proteins BBS2 and BBS7 were found to be missing in *D. melanogaster*, supporting previous studies that used different approaches such as reciprocal best BLAST (RBB) and Hidden Markov Model (HMM) searches (37,43). Interestingly, the only BBSome protein missing in *A. mellifera* was BBS1. Given that BBS1 is missing while other BBS1-like proteins are conserved in honeybees, it is likely that the function of BBS1 is now performed to some extent by BBS2, BBS7, or BBS9, which is also hypothesised for the *D. melanogaster* BBSome missing BBS2 and BBS7 (37). BBS1, BBS2, BBS7, and BBS9 all have similar protein architecture (N-terminal β-propeller, followed by an α-helical linker and ɣ-adaptin-ear domain); BBS1 differs in that it is missing the platform- and C-terminal α-helix domains present in BBS2, BBS7 and BBS9. Ultrastructure of the mammalian BBSome was recently established (18), which suggests that BBS1 mediates the interaction between ARL6 and the BBsome needed for membrane attachment. Interaction is facilitated by the ɣ-adaptin-ear domain of BBS1 but that domain is also present in the other ‘BBS1-like’ proteins, so BBS1 itself may be substitutable in the BBSome of some species. This is supported by the low sequence similarity of honeybee BBS1 to human BBS1 (Fig. 3A), which could result from relaxed purifying selection or positive selection acting in one or both species driving divergence.

**Fig. 3:**
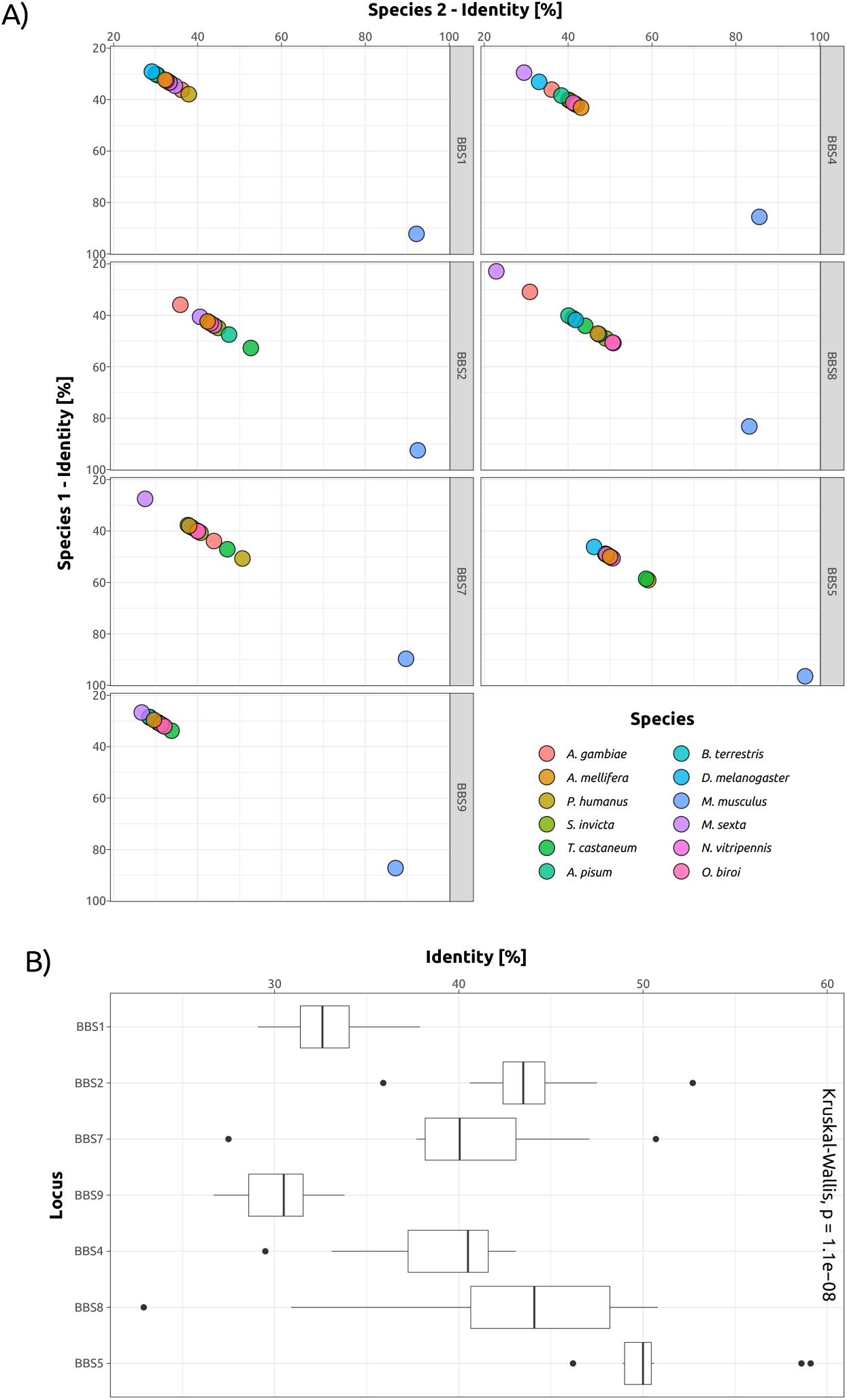
Variation in sequence similarity across insect BBS proteins. A) Scatterplots for each identified BBS protein across insect genomes displaying percentage identity at the protein level between human BBS proteins (species 1) and putative homologues in insects (n = 11) and mouse (species 2). Each species is represented by an individual colour. B) Boxplot displaying the distribution of percentage identify at the protein level for each BBS protein across all insect species examined (n = 11 species).

Overall, there was significant variation in terms of percentage identity shared between specific human BBS proteins and their insect homologues (min: 35% (BBS9); max: 54.8% (BBS5); Fig. 3A,B), which may reflect differences in terms of evolutionary conservation. While there is no single group of insects standing out in terms of conservation, there is a trend for individual BBS proteins to be generally more conserved than others. For example, the lowest sequence similarity to human BBS proteins is seen in BBS1 and BBS9, and highest in BBS5 (Fig. 3A). BBS8 has the most variation in sequence similarity of all BBSome proteins across insects (Fig. 3B). Low sequence similarities could reflect reduced constraints on functionality derived from specific amino acids, leaving room for evolutionary divergence and acquisition of novel functions. However, non-conserved structures may be amenable to loss, as could be the case for BBS1 in honeybees. The strikingly high sequence conservation of BBS5 on the other hand hints at potentially conserved functions, possibly outside cilia.

### BBS protein expression is different in tissues of queens, workers and males

To understand variation in gene expression profiles of BBS insect homologues, we examined differential gene expression of putative BBS homologues across sexes and tissues of *A. mellifera* (Supp. Table S2). Given that honeybee queens and males – referred to as drones – differ extensively in terms of physiology, morphology, and behaviour, yet share the same genome, such observed phenotypic differences are regulated via differential gene expression. Using datasets consisting of somatic and germline tissues from honeybee queens and males (49,50), we identified the expression of BBS4, BBS5, BBS7, and BBS8 homologues in *A. mellifera*, with BBS4 being the most highly expressed overall (Fig. 4A). While in terms of overall expression there were no significant differences between the sexes (Wilcoxon test, *p* > 0.05; Fig. 4B), we found significantly higher expression of both BBS5 and BBS8 in the brain compared to gonad (Wilcoxon test, *p*_*BBS5, brain-gonad*_ = 0.0001, *p*_*BBS8, brain-gonad*_ = 0.0451; Fig. 4C). This might reflect the high degree of specialisation insect cilia exhibit: in *D. melanogaster*, cilia are only expressed in (sensory) neurons and sperm cells. The fact that not all BBS genes are expressed higher in these tissues in honeybees is however intriguing and might hint at a more specialised role for these genes and their associated proteins, independent of cilia. We further compared the tissues of queens and drones (both within the same sex and versus the other sex; Fig. 4D) finding that BBS5 and BBS8 are significantly higher expressed in drone brains compared to drone gonads (Wilcoxon test, *p*_*BBS5, drone, brain-gonad*_ = *p*_*BBS8, drone, brain-gonad*_ = 0.0317; Benjamini-Hochberg corrected). The case is the same for BBS8 expressed in queen brains versus queen gonads (Wilcoxon test, *p*_*BBS8, queen, brain-gonad*_ = 0.0357; Benjamini-Hochberg corrected). When comparing expression in the brain, BBS8 is significantly lower expressed in queens compared to drones (Wilcoxon test, *p*_*BBS8, brain, queen-drone*_ = 0.0357; Benjamini-Hochberg corrected), and BBS5 is significantly higher expressed in queen gonads compared to drone gonads (Wilcoxon test, *p*_*BBS8, gonad, queen-drone*_ = 0.0357; Benjamini-Hochberg corrected). BBS5 and BBS8 both localise to mammalian nuclei (37) and both proteins have been shown to interact with E3-ubiquitin-protein ligase RING2 (RNF2) (29), a protein of the polycomb group (PcG) repressor complex 1 (PRC1). This complex is mainly responsible for histone 2A (H2A) monoubiquitylation (51,52), leading to repression of proteins crucial during development, such as those encoded by homeobox genes (53,54). Given the high degree of sequence conservation, this could also be the case in honeybees.

**Fig. 4:**
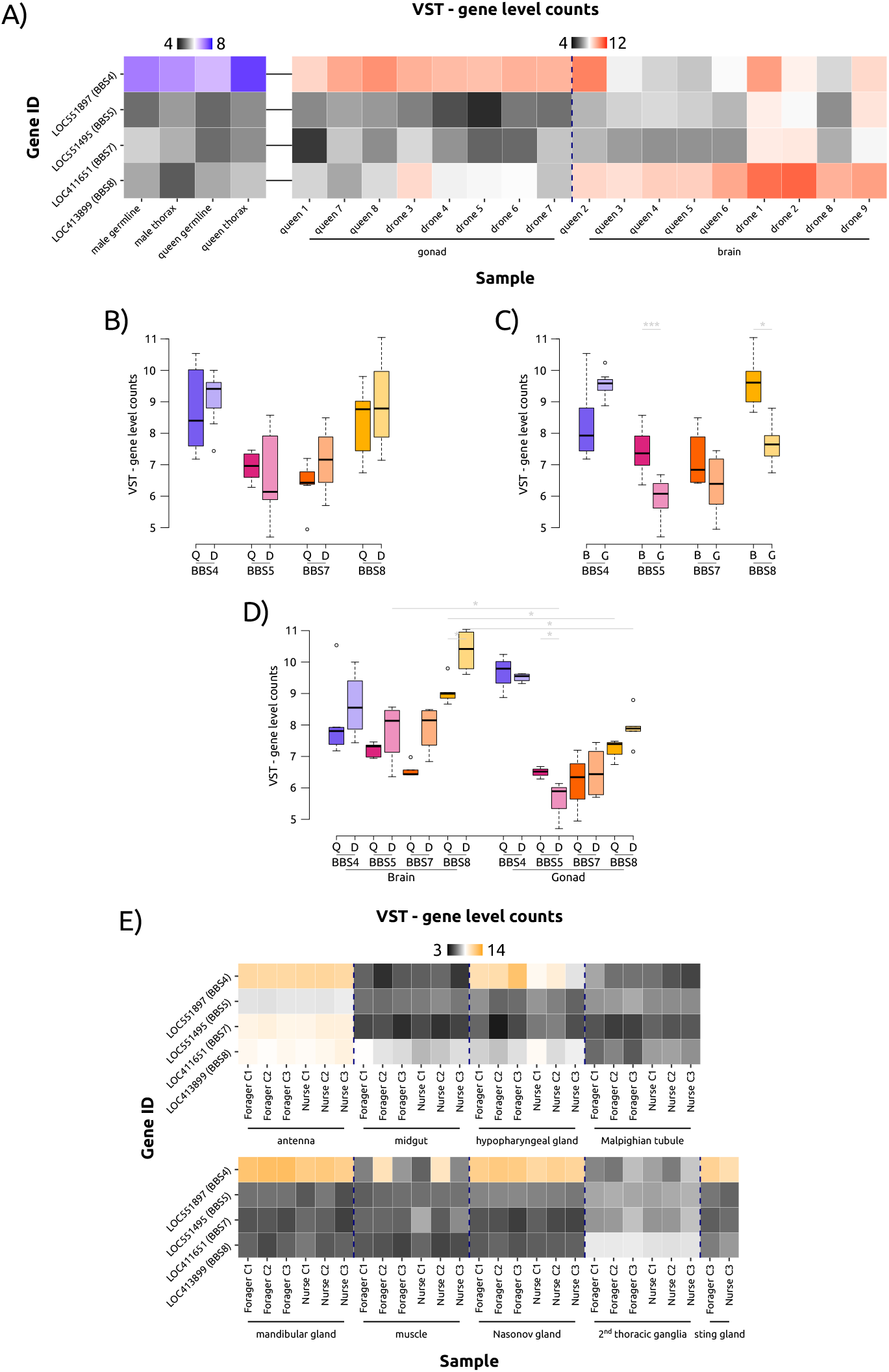
Expression differences of expressed honeybee BBS homologues from queens and drones, and across different tissues. A) Variance-stabilising transformed (VST) gene level counts of BBS proteins in somatic and germline tissues of honeybee queens and males. B) Expression profiles of BBS proteins in the brains and gonads of honeybee queens and drones. C) Expression profiles of BBS proteins across different tissues (Wilcoxon test, *: p <= 0.05; ***: p <= 0.001). D) Expression profiles of BBS proteins across tissues of different sexes (Wilcoxon test, *: p <= 0.05). E) VST gene level counts of BBS proteins in somatic tissues of honeybee foragers and nurse. Q: queen, D: drone; B: brain, G: gonad.

An alternative, larger tissue dataset consisting of different tissues from honeybee workers, and here especially from nurses and foragers, shows a similar picture: BBS4 is expressed quite frequently (and relatively strongly) in many tissue types, especially in glandular tissues and antennae (Fig. 4E). The consistently high expression of BBS4 in all honeybee glands studied, including the hypopharyngeal and mandibular glands in the head of the worker bees and the Nasonov and sting glands in their abdomen, suggests a specific function of this gene in glandular physiology, and possibly in chemical communication. Interestingly, the other BBS proteins are also relatively highly expressed in the antennae. Along with the eyes, the antennae are the most important sensory organs of the honeybee and have functions in chemosensory, tactile, temperature and acoustic perception. Type I and II cilia have been described for the Johnston organ of the honeybee antenna pedicel (55), and cilia have also been found in the odour-detecting sensilla placodea at the tip of the antennae (56). Higher expression of BBS proteins in antennae could therefore be due to their cilia function. To fully understand the possibly conserved role of BBS proteins in gene regulation, further studies of the potential target genes are required.

## CONCLUSION

The findings of this analysis predominantly validate earlier studies regarding the existence of BBS homologues in insects, such as *A. gambiae, A. mellifera*, and *D. melanogaster (43,44)*. While former studies focused on the evolution of molecular intraflagellar transport (44) and the evolutionary history of the centriole from protein components (43), our study investigated the evolutionary conservation and expression of BBS protein homologues across insect species. We were able to show that proteins forming the BBSome are potentially more conserved across insects than previously thought. Our study reveals that limiting homologue searches to few model organisms does not necessarily reflect the *status quo* across a larger set of species from the same family and might therefore skew conclusions drawn. In previous studies, homologues of all BBSome genes (with exception of BBS2 and BBS7) were described at the genomic level in *A. mellifera* and *D. melanogaster* (37,43,44). Interestingly, *A. mellifera* lacks a putative BBS1 homologue using our search criteria, despite being found in another study (43). This is seemingly a contradiction to previous work conducted on a Cape honeybee population (57), where the authors concluded LOC102655146, a locus annotated as BBS1-protein like, is under selective pressure for the social parasitism phenotype. Given that the protein architecture of BBS1 and BBS2 (also BBS7 and BBS9) are similar, it could be that Wallberg *et al*. erroneously ‘identified’ a BBS1-like protein as BBS1. It is still interesting to conduct further analyses towards the identified gene to shed light on possible altered protein interactions upon loss of one essential BBSome constituent.

The comparison between genomic/transcriptomic searches and proteomic approaches clearly shows that with seeds from distantly related species (human vs. insects in this case), orthology searches that focus on conservation of amino acid sequences are clearly superior, both in sensitivity and sensibility. Upon BLAST analysis, sequences of human BBS proteins were found both in the genome and in the transcriptome in each insect. The predicted expression of BBS8 in all but one queried species suggests that many insects transcribe BBS proteins, despite the absence of classical cilia in most tissues. This leads to the question which alternative function this BBS protein might perform in insects.

It is exciting to find a theoretically fully functional BBSome in insects despite their deviation from the ‘classical’ function of (primary) cilia throughout different tissues seen in other animals. Finding proteins differentially expressed between tissues in honeybees indicates that there are probably cilia-independent functions. The differential expression of BBSome proteins in ciliated tissues, such as the brains of queens and drones, suggests that these proteins may have additional functions beyond their role in ciliary transport.

## MATERIALS AND METHODS

### Data procurement

The data used for BLAST analyses of transcriptomes and genomes was obtained from the National Center for Biotechnology Information nucleotide database. For each insect, the representative genome and transcriptome GenBank assemblies were used for this purpose. A list of the GenBank assembly accessions can be found in Supp. Table S3. The human BBS accession IDs used as a seed can be found in Supp. Table S4.

### BLAST analysis and homologue search

In terms of BLAST analysis (59) a hit is defined by a detected significant similarity between the sequences compared with respect to the chosen parameter values. Hits can differ in aspects like position, length, e-value and score. In this study the e-value was chosen as representation value to classify the significance and display hits. The BLASTn analyses were carried out via the NCBI BLAST online tool (general parameters: expect threshold = 0.05, word size = 11; scoring parameters: match/mismatch scores= 2,-3, gap costs = existence: 5 extension: 2).

To determine homology of BBS-associated proteins across diverse insect orders, predicted RefSeq proteomes were downloaded from NCBI for the following species: Diptera, *Drosophila melanogaster, Anopheles gambiae*; Hymenoptera, *Apis mellifera, Bombus terrestris, Nasonia vitripennis, Solenopsis invicta, Ooceraea biroi*; Lepidoptera, *Manduca sexta*; Hemiptera, *Acyrthosiphon pisum*; Phthiraptera, *Pediculus humanus*; Coleoptera, *Tribolium castaneum*; and Mammalia, *Homo sapiens* and *Mus musculus*. From each, the longest predicted peptide was extracted using the OrthoFinder (v.2.5.4) script primary_transcript.py (48). OrthoFinder was subsequently used (default settings) to perform pairwise alignments between each species allowing for the identification and generation of orthogroups, which are groups consisting of potential homologues across species of interest. Such orthogroups were parsed for the determination of insect homologues of described human BBS-associated genes.

### Differential gene expression analysis

For the purposes of examining gene expression of putative BBS-associated homologues, we examined expression profiles in somatic and germline tissues of the Western honeybee, *A. mellifera (60)*, BioProject ID: PRJNA243651; (49), BioProject ID: PRJNA386859; (50), BioProject ID: PRJNA689223). We downloaded publicly available transcriptomic datasets from the NCBI Short Read Archive using the sra-toolkit. For each library, we extracted the data in FASTQ format and performed quality assessments using FastQC v.0.11.9 (https://www.bioinformatics.babraham.ac.uk/projects/fastqc/). Data were filtered using fastp v.0.23.2 (61) to remove low quality reads and trim adaptors. We then pseudoaligned each sample against an indexed predicted transcriptome using kallisto v.0.44.0 (62) providing a transcript-level quantification of expression. Using these estimates, we generated gene-level counts using tximport v1.26.1 (63) and for each dataset, generated variance-stabilised transformed data using DESeq2 v1.38.3 (64).

### Statistical testing

Datasets used for statistical testing can be found in Supp. Table S2. Tests were performed in R v.4.0.3 (65), with packages tidyverse (66) (including dplyr (67), ggplot2 (68) and stringr (69)), reshape2 (70), and FSA (71). The accompanying RMarkdown notebook is available figshare (https://doi.org/10.6084/m9.figshare.23780871). Datasets were tested for normality by Shapiro-Wilk testing (where normality was assumed if p > 0.05) and depending on outcome, different tests were used to compare subsets. Comparisons were done by Wilcoxon rank-sum test (p > 0.05: not significant; 0.05 >= p > 0.01: *; 0.01 >= p > 0.001: **; 0.001 >= p > 0.0001: ***). Multiple testing correction was done using the Benjamini-Hochberg procedure (72).

### Additional software

Figures were prepared with ggpubr (73) and Inkscape (74).

## ACKNOWLEDGEMENTS

This project was funded by the Deutsche Forschungsgemeinschaft (DFG, German Research Foundation) – GRK2526/1 – Projectnr. 407023052.

